# Flexible control of Pavlovian-instrumental transfer based on expected reward value

**DOI:** 10.1101/2021.04.08.438512

**Authors:** Andrew T. Marshall, Briac Halbout, Christy N. Munson, Collin Hutson, Sean B. Ostlund

**Author notes:** Correspondence Dr Sean Ostlund, Department of Anesthesiology & Perioperative Care, University of California, Irvine School of Medicine, Irvine, CA 92697, USA.

## Abstract

The Pavlovian-instrumental transfer (PIT) paradigm is widely used to assay the motivational influence of reward-predictive cues, reflected by their ability to invigorate instrumental behavior. Leading theories assume that a cue’s motivational properties are tied to predicted reward value. We outline an alternative view which recognizes that reward-predictive cues may suppress rather than motivate instrumental behavior under certain conditions, an effect termed positive conditioned suppression. We posit that cues signaling imminent reward delivery tend to inhibit instrumental behavior, which is exploratory by nature, in order to facilitate efficient retrieval of the expected reward. According to this view, the motivation to engage in instrumental behavior during a cue should be *inversely* related to the value of the predicted reward, since there is more to lose by failing to secure a high-value reward than a low-value reward. We tested this hypothesis in rats using a PIT protocol known to induce positive conditioned suppression. In Experiment 1, cues signaling different reward magnitudes elicited distinct response patterns. Whereas the 1-pellet cue increased instrumental behavior, cues signaling 3 or 9 pellets suppressed instrumental behavior and elicited high levels of food-port activity. Experiment 2 found that reward-predictive cues suppressed instrumental behavior and increased food-port activity in a flexible manner that was disrupted by post-training reward devaluation. Further analyses suggest that these findings were not driven by overt competition between the instrumental and food-port responses. We discuss how the PIT task may provide a useful tool for studying cognitive control over cue-motivated behavior in rodents.

---

Reward-paired cues acquire Pavlovian incentive motivational properties which allow them to invigorate instrumental reward-seeking behavior (Dickinson et al. 2000; Estes 1948; Lovibond 1981), a phenomenon referred to as Pavlovian-instrumental transfer (PIT). This influence seems to serve an adaptive function by promoting risky and effortful foraging activity in environments that signal potential reward availability. The PIT paradigm is widely used to study the mechanisms of cue-motivated behavior (Cartoni et al. 2016; Corbit and Balleine 2016) and how they contribute to pathological reward seeking in addiction and related disorders (Corbit and Janak 2007; 2016; Garbusow et al. 2016; Genauck et al. 2020; LeBlanc et al. 2013; LeBlanc et al. 2014; LeBlanc et al. 2012; Marshall and Ostlund 2018; Ostlund et al. 2014; Saddoris et al. 2011; Sebold et al. 2021; Shiflett 2012; Shiflett et al. 2013; Vogel et al. 2018; Wyvell and Berridge 2001). However, despite decades of research, much remains unclear about how fundamental variables such as expected reward value influence expression of the PIT effect.

Leading computational models of incentive learning (Dayan and Balleine 2002; McClure et al. 2003; Zhang et al. 2009) assume that motivational value is assigned to cues based on the total amount of delay-discounted reward that they predict (i.e., their state value). The motivational influence of cues should therefore directly depend on basic Pavlovian conditioning parameters such as reward probability, cue-reward interval, and reward magnitude. This account makes some intuitive predictions. For instance, a cue that reliably signals the immediate delivery of a large reward should acquire strong motivational properties, whereas a cue signaling that an upcoming reward will be small, delayed, or unlikely to occur at all, should acquire weak motivational properties.

However, in contrast to these predictions, evidence suggests that the motivational impact of reward-paired cues is instead *inversely* related to their ability to predict reward. For instance, cues signaling a high probability of imminent reward do not invigorate and may even suppress instrumental performance, an effect known as positive conditioned suppression (Azrin and Hake 1969; Crombag et al. 2008; Lovibond 1981; Meltzer and Hamm 1978; Miczek and Grossman 1971; Vandyne 1971). Instead, such cues elicit high levels of Pavlovian conditioned approach behavior directed at the food-port (Marshall et al. 2020). In contrast, cues that signal a low probability of reward become potent motivators of instrumental behavior, while eliciting more modest levels of food-port activity (Marshall et al. 2020). The motivational influence of cues also depends on their temporal relationship with reward delivery (Delamater and Holland 2008; Delamater and Oakeshott 2007; Lovibond 1981; Matell and Della Valle 2018). For instance, we have shown that cues signaling a fixed 30-sec interval between cue onset and reward delivery produce a gradual suppression of instrumental behavior and a coincident increase in food-port activity as the expected reward delivery time draws near (Marshall and Ostlund 2018).

Such findings suggest that reward-paired cues can acquire distinct motivational and predictive properties that evoke different kinds of behavior, with the former promoting the pursuit of new rewards through instrumental behavior and the latter eliciting a pause in instrumental behavior and anticipatory reward-retrieval activity (Ostlund and Marshall 2021). Organizing behavior in this manner, based on reward expectancy, is important for efficient foraging and is a central pillar of behavior systems theory (Timberlake et al. 1982). While it is adaptive to seek out new rewards through instrumental behavior (or other general search activities) when rewards are scarce, such behavior is unnecessary and may even interfere with the retrieval of a reward that is expected soon (i.e., focal search), increasing the chance that it will be pilfered or otherwise lost.

However, this simplistic description of foraging sidesteps the complexity involved in ambiguous situations, when cues may elicit conflicting tendencies to both seek out new rewards *and* collect an expected reward. The factors involved in resolving such conflict are not well understood, though presumably the value of the expected reward plays an important role. Early studies on the interaction between Pavlovian and instrumental learning processes may also provide some insight. For instance, Konorski and colleagues demonstrated through a series of studies that Pavlovian cues predicting imminent reward do not simply elicit conditioned consummatory responses – in their case, salivation and orienting toward the food cup – they also acquire the ability to acutely interrupt ongoing instrumental behavior (Ellison and Konorski 1965; Konorski 1967). Such cues were also able to prevent other discriminative cues from motivating instrumental performance (Soltysik et al. 1976) and were themselves extremely resistant to acquiring motivational properties if later used as discriminative stimuli for instrumental performance (Konorski and Wyrwicka 1950). These findings suggest that cues signaling imminent, response-independent reward actively inhibit the expression of cue-motivated behavior.

Following this logic, we have hypothesized that Pavlovian cues acquire the potential to motivate instrumental behavior, but that this motivational response is subject to cognitive control and is therefore suppressed in situations where such behavior would be disadvantageous (Marshall et al. 2020; Marshall and Ostlund 2018; Ostlund and Marshall 2021). Cognitive control broadly refers to the ability to regulate thoughts, emotions, and actions based on an internally represented behavioral goal (Braver 2012). A core function of cognitive control is to override prepotent response tendencies – particularly those which are impulsive or habitual in nature – in situations where an alternative course of action is more advantageous (Botvinick and Braver 2015; Miller and Cohen 2001). We can apply this framework to PIT by assuming that when a cue signals that a valuable reward is imminent, the impulse to engage in instrumental reward-seeking behavior will be suppressed in order to allow for efficient reward retrieval. Thus, the value of the reward predicted by a cue should play an important albeit indirect role in modulating instrumental performance, indicating the degree to which such behavior should be suppressed. A cue that signals imminent delivery of a high-value reward should therefore be less effective in motivating instrumental behavior and more effective in eliciting food-port activity than a cue that signals a less valuable reward. These predictions are readily distinguished from those made by theories of incentive learning that assume a positive correlation between motivational and predictive value (Dayan and Balleine 2002; McClure et al. 2003; Zhang et al. 2009).

Interesting, the limited data that exist on this subject suggests that expected reward value rarely affects the expression of PIT performance. For instance, several studies have shown that cues retain their ability to motivate instrumental behavior despite predicting a reward that has been devalued through conditioned taste aversion learning (Colwill and Rescorla 1990; Holland 2004; Rescorla 1994). However, such studies have generally used PIT protocols designed to maximize motivational effects and use cues that signal long, sporadic intervals before reward delivery. Therefore, while these findings suggest that cues normally motivate instrumental behavior independently of expected reward value, they do not address whether expected reward value modulates the suppressive influence of cues that signal imminent reward (i.e., positive conditioned suppression).

We investigated this issue using an alternative version of the PIT task in which reward-predictive cues tend to suppress instrumental behavior and increase food-port activity in apparent anticipation of reward delivery (Marshall and Ostlund 2018). The role of expected reward value on these responses was assessed using a parametric manipulation of reward magnitude in Experiment 1 and post-training reward devaluation in Experiment 2.

## Experiment 1

Experiment 1 investigated the influence of expected reward magnitude on the expression of instrumental reward-seeking and food-port entry behavior during PIT testing. Briefly, hungry rats were first trained to lever press for food pellets before undergoing Pavlovian conditioning, in which two distinct 30-sec auditory cues signaled food pellet delivery at cue offset. Reward magnitude was varied across cues and groups. While all groups had at least one cue that signaled three food pellets, the alternate cue signaled either one (Group 1/3), three (Group 3/3), or nine food pellets (Group 3/9). PIT testing was then performed by intermittently presenting these reward-predictive cues in a noncontingent manner while rats were free to press the lever and enter the food-port, in the absence of reward delivery.

## Methods

### Animals and Apparatus

Twenty-six experimentally naïve adult male Long Evans rats (Envigo) were used in this experiment. They arrived at the facility (University of California, Irvine; Irvine, CA, USA) at approximately 10 weeks of age. They were pair-housed in a colony room set to a standard 12:12 hr light:dark schedule. Rats were tested during the light phase. Water was always provided ad libitum in the home cages. Rats were fed between 10-14 g of standard lab chow per day during the experiment to maintain them at ∼85% of their estimated free-feeding bodyweight. Husbandry and experimental procedures were approved by the UC Irvine Institutional Animal Care and Use Committee (IACUC) and conducted in accordance with the National Research Council Guide for the Care and Use of Laboratory Animals.

The experiment was conducted in 14 operant chambers (Med-Associates; St. Albans, VT), each housed within sound-attenuating, ventilated boxes. Each chamber was equipped with a stainless-steel grid floor; two stainless steel walls (front and back); and a transparent polycarbonate side-wall, ceiling, and door. Pellet dispensers, mounted on the outside of the operant chamber, were equipped to deliver 45-mg food pellets (Bio-Serv) into a recessed food-port centered on the lower section of the front wall. Head entries into the food-port were transduced by an infrared photobeam. A retractable lever was located to the left of the food-port, on the front wall. The chamber was also equipped with a house light centered at the top of the back wall. Auditory stimuli were presented to animals via a speaker located on the back wall. Experimental events were controlled and recorded with 10-ms resolution by the software program MED-PC IV (Tatham and Zurn 1989).

### Procedure

#### Magazine training

All sessions of all phases began with the onset of the houselight. In each of two 30-minute sessions of magazine training, grain-based food pellets were delivered on a random-time (RT) 60-s schedule.

#### Instrumental training

During initial instrumental (lever-press) training, rats were continuously reinforced with a grain-based food pellet delivery for pressing the left lever (fixed-ratio, FR-1), earning a maximum of 30 pellets per session. These FR-1 sessions lasted no more than 30 min. Seven rats required an extra session of FR-1 training, which lasted until these rats earned 30 pellets. During subsequent training sessions, lever pressing was reinforced according to a random-interval (RI) schedule, such that the lever remained available but was inactive for an average of *t* seconds after each reward delivery, where individual *t* values were selected randomly from an exponential distribution. The RI schedule was changed over training days with 1 day of RI-5 (*t* = 5 sec), 1 day of RI-15 (*t* = 15 sec), 2 days of RI-30 (*t* = 30 sec), and 10 days of RI-45 (*t* = 45 sec) training. Each RI session lasted 30 minutes.

#### Pavlovian training

Pavlovian training involved exposure to two 30-s conditioned stimuli (CS; 3-kHz tone and 10-Hz clicker; 80 dB) which were paired with reward (grain-based food pellets). Rats were assigned to one of three groups with different CS-reward magnitude arrangements. For Group 1/3 (*n* = 9), one CS terminated with 1 pellet and the other with 3 pellets. For Group 3/3 (*n* = 8), both CSs terminated with 3 pellets. For Group 3/9 (*n* = 9), one CS terminated with 3 pellets and the other with 9 pellets. Stimulus identity was counterbalanced with group and reward magnitude conditions.

In each 20-min session, a 60-s interval preceded onset of the first CS. There was a random 120-s inter-trial-interval (ITI) between CS presentations, and a 60-s interval following the final CS presentation prior to the end of the session. Pavlovian training lasted for 25 daily sessions, each involving 4 pseudorandomly-alternating presentations of each CS (8 total trials per session).

#### Pavlovian-to-instrumental transfer (PIT)

Following Pavlovian training, rats received two daily sessions of instrumental retraining (identical to earlier sessions with the RI-45 schedule) followed, on the next day, by one 30-min session of instrumental extinction, in which the lever was continuously available but was inactive. On the following day, rats received a PIT test session (30.25 minutes in duration), during which the lever was once again continuously available but inactive. During the test, rats received 4 noncontingent presentations of each 30-s CS in pseudorandom order (ABBABAAB). The ITI was 150s, and a 5-min interval preceded onset of the first CS. No food pellets were delivered at test.

### Data Analysis

All summary measures were obtained from the raw data using MATLAB (The MathWorks; Natick, MA, USA), and analyzed with mixed-effects regression models (Pinheiro and Bates 2000), a powerful analytical framework that is both well established and highly recommended for behavioral research (Boisgontier and Cheval 2016). Mixed-effects models are comparable to repeated-measures regression analyses, and allow for parameter estimation per manipulation condition (fixed effects) and the individual (random effects) (Bolker et al. 2009; Hoffman and Rovine 2007; Pinheiro and Bates 2000; Schielzeth et al. 2013). Mixed-effects regression models (1) effectively handle missing data and (2) permit the inclusion of categorical and continuous predictors in the same analysis, thus allowing detection of group-level changes across ordered data samples (i.e., continuous time points) while also accounting for corresponding individual differences. All relevant fixed-effects factors were included in each model. Categorical predictors were effects-coded (i.e., codes sum to 0), and continuous predictors were mean-centered. For analyses of Pavlovian training and PIT, the fixed-effects structure included main effects of group and reward magnitude, and the random-effects structure included a by-rat random intercept. PIT analyses also included main effects of time within the CS (i.e., CS 10-s period), with CS Reward Magnitude × CS 10-s Period interactions also included for analysis of individual groups. (Here, analysis of Group 1/3’s food-port-entry behavior employed a linear model with no random effects due to random-effects convergence issues given outlier removal, described below.) Both group (categorical) and reward magnitude (continuous; i.e., 1, 3, 9) were included in these analyses to differentiate overall between-group differences in behavior versus sample-wide changes in behavior as a function of differences in reward magnitude. Instrumental training analyses incorporated generalized linear mixed-effects models (family: gamma, link: log) with predictors of group and time since the previous reward delivery. The random-effects structure included a random by-rat slope of time since reward delivery and a random intercept of rat, restricted to be uncorrelated. The alpha level for all tests was .05. Sample sizes were not predetermined based on statistical analysis but are similar to those reported in previous publications (Halbout et al. 2019; Marshall et al. 2020; Marshall and Ostlund 2018). Main effects and interactions are reported as the results of ANOVA *F*-tests (i.e., whether the coefficients for each fixed effect were significantly different from 0).

Our primary dependent measures were the rates of lever pressing and food-port entry behavior (recorded as the number of discrete food-port beam breaks). We quantified cue-induced changes in behavior by subtracting the mean response rate (response per minute) during pre-CS periods (30 sec each) from the mean response rate during CS periods, calculated separately for consecutive 10-sec periods within individual 30-sec CSs to characterize the time course of responding. Pre-CS (baseline) data were averaged across all CS trials (within-subject).

The final five sessions of Pavlovian training were used to assess conditioned food-port entry behavior during CS trials relative to pre-CS baseline periods. Analyses of instrumental training included the final three sessions of training. Data points were considered outliers if their values were at least three scaled median absolute deviations from the median (Leys et al. 2013), in which the median value of the absolute deviations from the median was scaled by ∼1.48 (Rousseeuw and Croux 1993) and then multiplied by 3; for PIT analyses, outliers were based on the rats’ mean difference scores within each condition. For the current experiment, 24 individual data points were removed from the instrumental training analysis (i.e., 24 of 1,170 data points [26 rats × 45 bins]) and one rats’ data (Group 1/3, 3-pellet CS) were removed from the PIT analyses of food-port-entry behavior. Lastly, 38 of 528 data points (i.e., paired observations of lever-pressing and food-port-entry behavior) were excluded from trial-by-trial analyses of concurrent local changes in both CS-induced lever-pressing and food-port-entry behavior. There were no outliers in the other analyses. Outlier removal was specific to each analysis, such that a rats’ exclusion from one analysis did not prohibit inclusion from other analyses.

## Results

### Instrumental and Pavlovian Training

Rats were first trained to lever press for food reward. While the rats had not yet been assigned to groups for Pavlovian training, Figure 1A shows the groups’ mean lever-press rates on an RI-45 s schedule of reinforcement as a function of time since the previous reward delivery. All groups [mean (SEM)] lever pressed at comparable rates [Group 1/3: 15.8 (1.8); Group 3/3: 14.8 (1.4); Group 3/9: 15.7 (1.5)]. Per a generalized linear mixed-effects model (distribution = gamma, link = log) on response rates during the 15-45 s time window, there was neither a main effect of group, *F*(1, 786) = 0.21, *p* = .812, nor a Group × Time interaction, *F*(2, 786) = 0.77, *p* = .461.

**Figure 1.**
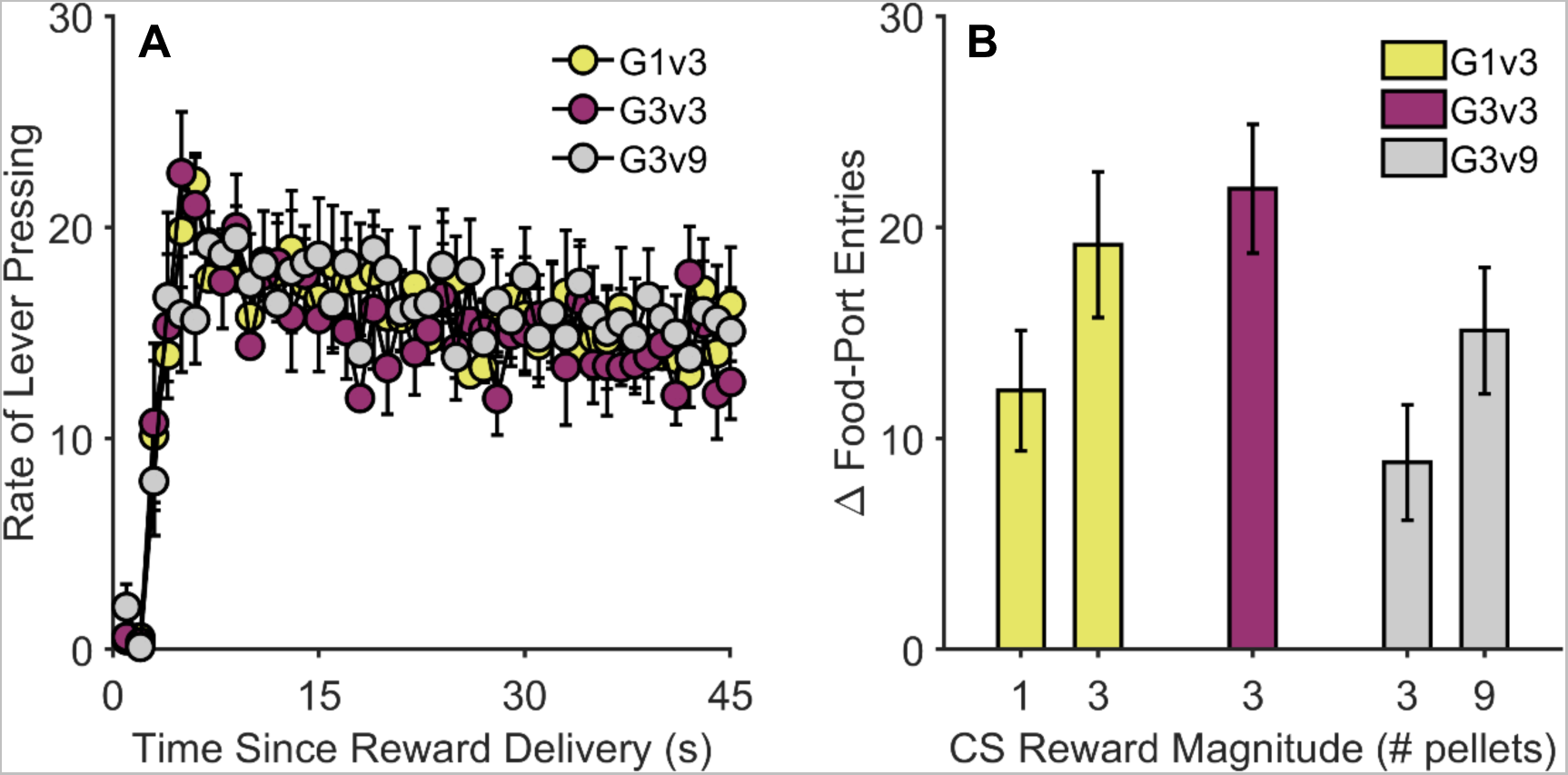
Instrumental and Pavlovian training in Experiment 1. (A) Lever-press rates on a random-interval (RI) 45-s schedule of reinforcement as a function of time (in seconds) since previous reward delivery. (B) CS-induced increases in food-port entry rate relative to the pre-CS baseline. CS = conditioned stimulus. G1v3 = Group 1/3. G3v3 = Group 3/3. G3v9 = Group 3/9. Error bars reflect ± 1 between-subjects standard error of the mean.

Following instrumental training, rats were trained to associate two 30-s CSs with food reward, delivered upon termination of the cues. Figure 1B shows conditioned food-port entry behavior (relative to the pre-CS baseline) for the final 5 sessions of training. Analysis of these data revealed a main effect of group, *F*(2, 40) = 5.69, *p* = .007, and a main effect of CS reward magnitude, *F*(1, 40) = 16.27, *p* < .001. The latter effect appeared to be driven by Groups 1/3 and 3/9, which showed higher levels of food-port activity during whichever CS signaled the larger of the two possible reward magnitudes.

### Pavlovian-instrumental transfer (PIT)

Following two sessions of instrumental retraining and one session of instrumental extinction, rats were given a PIT test, in which the 30-s CSs were presented while the rats were able to freely lever press without reinforcement. Figures 2A and 2B show rats’ CS-induced change in lever-press rate and food-port entry rate, respectively, relative to pre-CS baseline periods. Analyses revealed main effects of group, *F*(2, 127) = 3.59, *p* = .030, and CS reward magnitude on lever pressing, *F*(1, 127) = 5.74, *p* = .018, in which CS-induced lever pressing decreased with increases in predicted reward magnitude. For food-port entry rate, there was a main effect of group, *F*(2, 124) = 4.71, *p* = .011, and a significant increase in CS-induced food-port entries with predicted reward magnitude, *F*(1, 124) = 4.45, *p* = .037.

**Figure 2.**
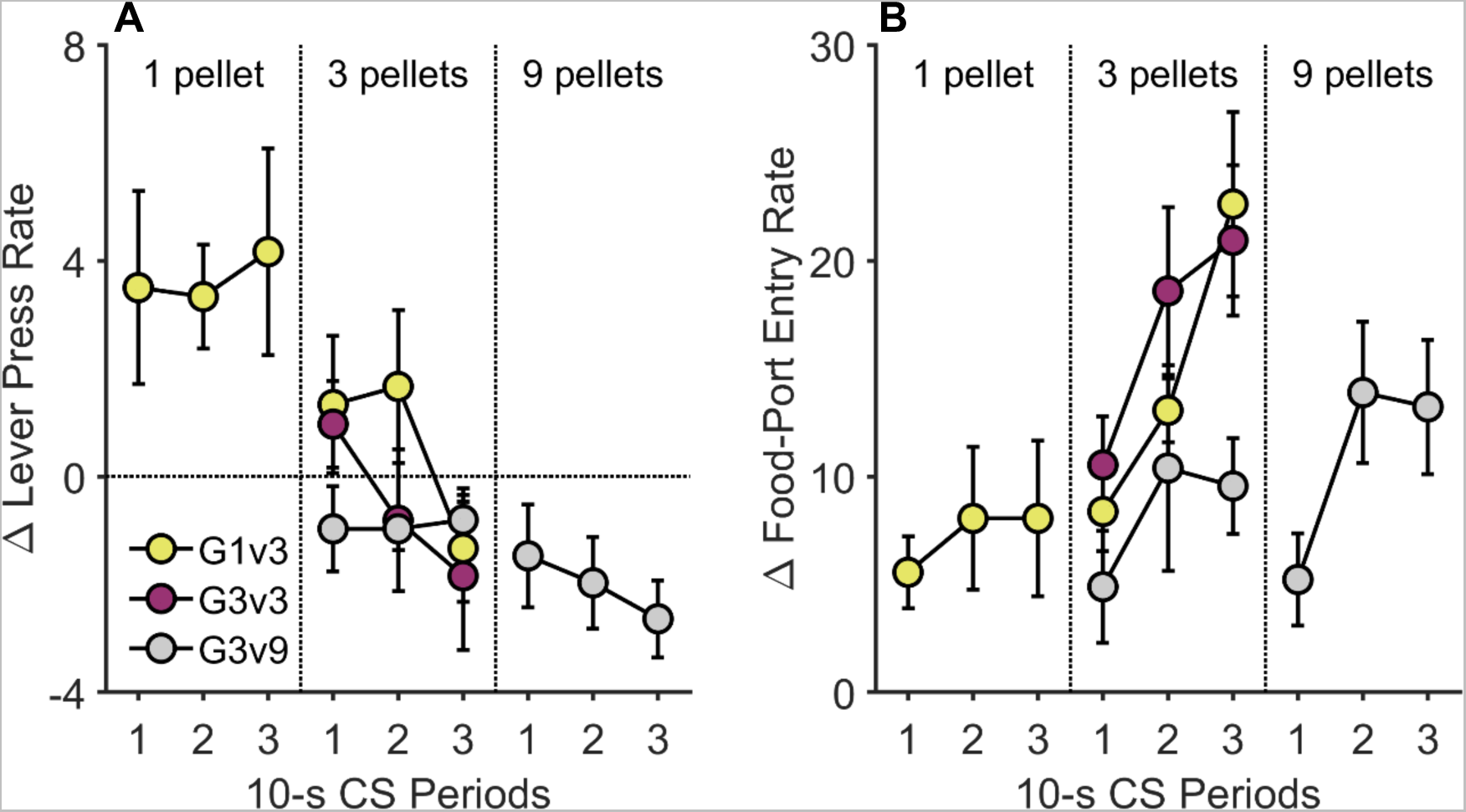
Pavlovian-instrumental transfer in Experiment 1. CS-induced changes in (A) lever-press rates and (B) food-port entries (relative to the pre-CS baseline) on the Pavlovian-instrumental transfer (PIT) test. These data are plotted separately according to Group and Reward Magnitude and show changes in responding over time (10-sec bins) during 30-sec CS presentations (averaged across trials). CS = conditioned stimulus. G1v3 = Group 1/3. G3v3 = Group 3/3. G3v9 = Group 3/9. Error bars reflect ± 1 between-subjects standard error of the mean.

Given the between-groups manipulation of reward magnitude and that one group experienced only one reward magnitude (i.e., Group 3/3), a second set of within-group analyses were conducted to better elucidate how differential reward magnitude influenced CS-induced changes in behavior. For Group 1/3, the 3-pellet CS reduced press rates more than the 1-pellet CS, *F*(1, 50) = 8.37, *p* = .006, and also led to higher levels of food-port entry, *F*(1, 47) = 10.08, *p* = .003. The food-port entry analysis also detected a significant CS Reward Magnitude × CS 10-s Period interaction, *F*(1, 47) = 4.16, *p* = .047, which indicated that the increase in food-port entries over time was steeper for the 3-pellet CS than the 1-pellet CS. Similarly, for Group 3/9, lever pressing was suppressed more by the 9-pellet CS than the 3-pellet CS, *F*(1, 50) = 5.36, *p* = .025, but there was no effect of reward magnitude on food-port entry, *F*(1, 50) = 1.79, *p* = .187.

These findings are consistent with the cognitive control hypothesis but could be the product of response competition between food-port and lever-press behaviors. Specifically, when given a cue that signals a large, desirable reward, rats may simply lose the opportunity to lever press because they are preoccupied with checking the food-port. However, we found little support this alternative account. For instance, if a rat’s tendency to check the food-port during a specific cue period interferes with their ability to also press the lever during that period, then these responses should be negatively correlated across cue periods for individual rats (e.g., large increases in food-port activity should co-occur unchanged or decreased press rates). To investigate this possibility, we assessed the within-subject correlation between press- and entry-rate difference scores across all 10-sec CS time bins (3 bins per trial x 4 trials, as in Figure 2), separately for each CS type, for each rat. Figure 3 shows that, with the exception of one CS type for each of 2 rats, *r*’s ≤ -0.59, *p*’s ≤ .045, lever pressing and food-port entries were *not* significantly correlated, *p*’s ≥ .050 (median *p*-value = .467). Moreover, of the 2 rats that did show significant correlations between press and entry rate, neither correlation passed Bonferroni correction for multiple comparisons (i.e., .05 divided by 44 separate correlations = .0011). Also notable was the finding that rats were in many cases able to increase their rate of lever pressing (i.e., displaying positive difference scores) during cue periods with extreme increases food-port entry behavior (> 30 entries/min), suggesting that food-port activity did not impose a major limit on lever-press performance.

**Figure 3.**
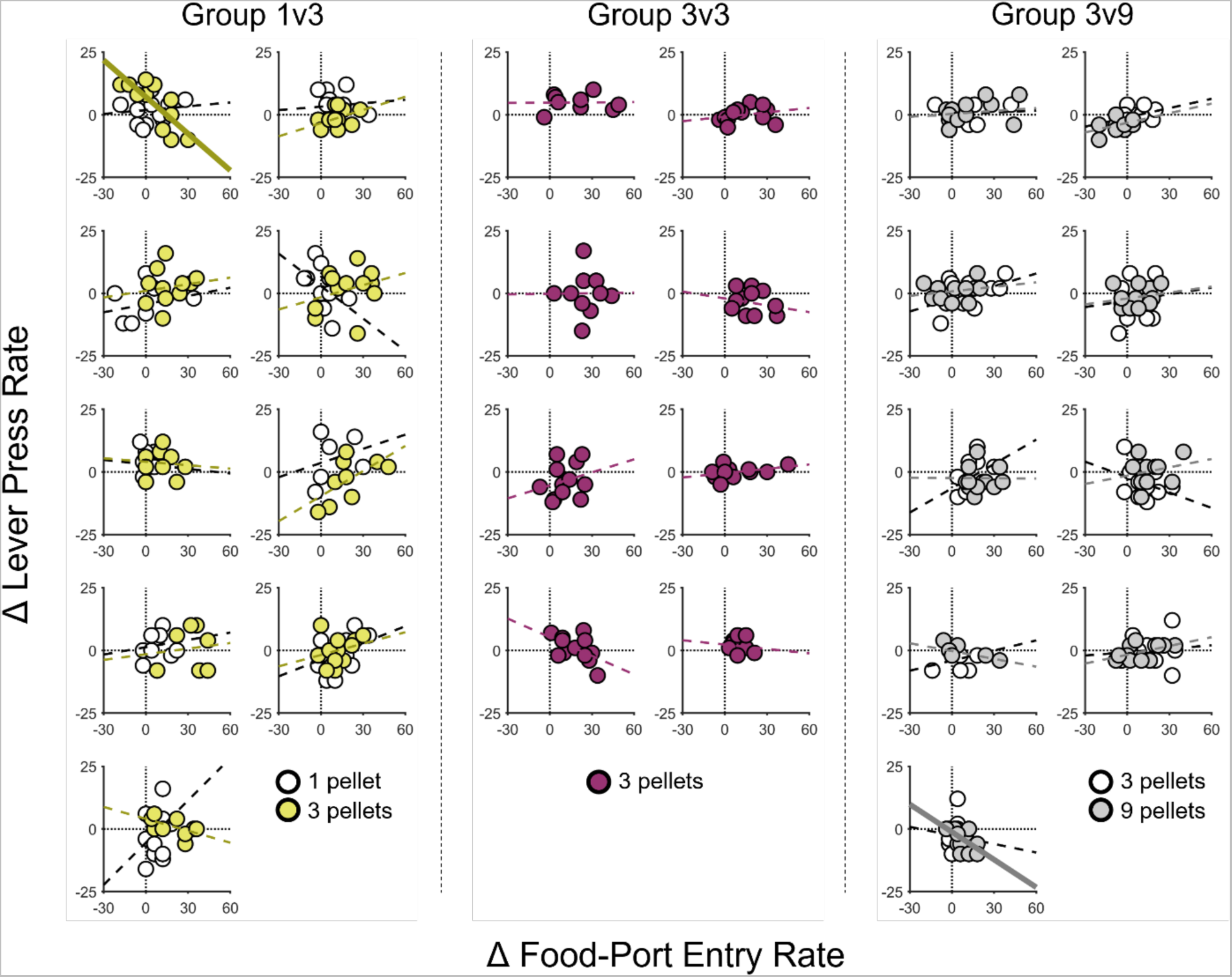
Trial-by-trial relationship between food-port-entry and lever-press rate for each 10-s CS bin during the Pavlovian-instrumental transfer (PIT) test for individual rats in Experiment 1. Scatterplots are presented for each rat. Data are separated by trial (unlike in Figure 2) and 10-s CS bin (as in Figure 2). The abscissa shows CS-induced changes in food-port-entry rate for individual time bins; the ordinate shows CS-induced changes in lever-press rate for corresponding time bins. For each CS type for each rat, simple regression lines were fit, with dashed lines indicating that the fit was not significant and bolded solid lines indicating significance at *p* < .05. CS = conditioned stimulus.

We also assessed whether food-port and lever-press responses competed more acutely over short time frames. For this analysis, we constructed peri-event histograms showing the probability of food-port entry occurrence during 0.1-sec periods surrounding individual lever-press responses (Figure 4A-C) and the corresponding probability of lever-press occurrence surrounding individual food-port entry responses (Figure 4D-F); a +/- 8-sec peri-event window was used for the panels in Figure 4. Visual inspection of these data indicates that there were only very brief (∼ 1-sec) dips in the probability of food-port behavior when rats lever pressed (Figures 4A-C) and in the probability of lever-pressing when rats entered the food-port (Figures 4D-F). Importantly, we also observed pronounced *increases* in the probability of food-port checking shortly after lever-press performance (Figures 4A-C) and in the probability of lever-press prior to food-port entry behavior (Figures 4D-F). These findings are consistent with previous reports (Halbout et al. 2019; Marshall et al. 2020; Marshall and Ostlund 2018) and indicate that these responses were typically performed together as a lever-pressàport-entry action sequence.

**Figure 4.**
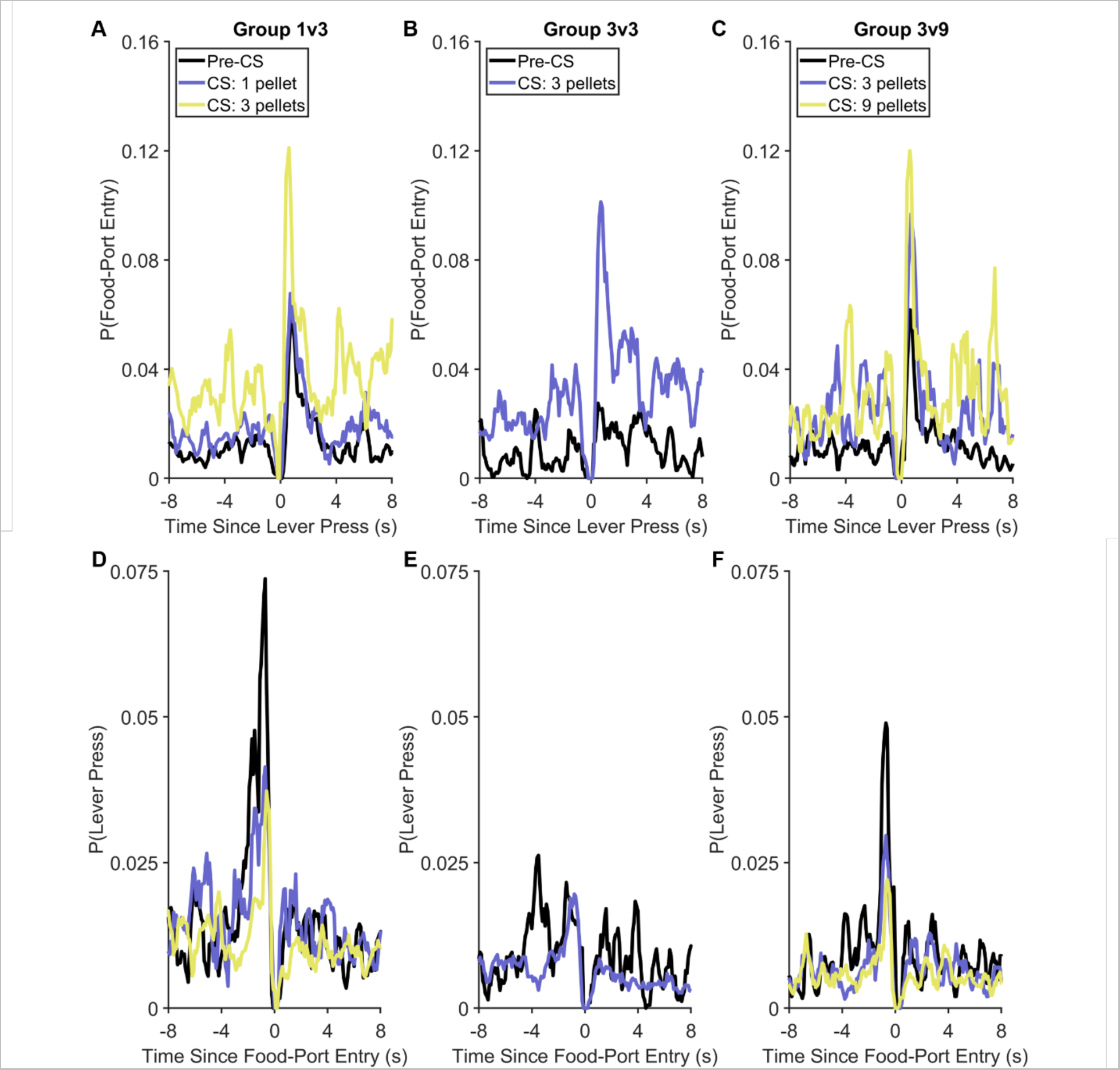
Peri-event histograms for lever-pressing and food-port-entry behavior during the Pavlovian-instrumental transfer test in Experiment 1. **(A-C)** Probability of food-port entries as a function of time from each lever press. **(D-F)** Probability of lever presses as a function of time from each food-port entry. Note the y-axes are different for Panels A-C and Panels D-F.

## Discussion

The results of Experiment 1 indicate that the motivational influence of reward-predictive cues on instrumental behavior varies inversely with expected reward magnitude, with the 1-pellet cue increasing reward seeking and the 3- and 9-pellet cues decreasing this behavior, particularly near the expected time of reward delivery. In contrast, attempts to retrieve reward at the food-port were more apparent when larger rewards were expected. While we did not include a control for nonassociative effects (e.g., pseudoconditioning) in this study, we have included such controls in several previous studies using similar designs and typically find that unpaired cues have little or no effect on instrumental performance (LeBlanc et al. 2013; LeBlanc et al. 2014; Ostlund et al. 2014). Moreover, the parametric effects of reward magnitude strongly imply that CS-induced changes in lever-press rate were attributable to CS-specific associative learning.

It is also notable that cue-elicited food-port entries did not monotonically vary with the magnitude of the predicted reward during either Pavlovian training or PIT testing, particularly in Group 3/9. While this may reflect random variability, our informal observations suggest that cues signaling larger rewards tend to elicit fewer but more persistent bouts of waiting at the food-port than the other cues. We therefore recorded and analyzed food-port dwell times in Experiment 2 to more fully characterize the influence of predicted reward value on this behavior and further probe competition between food-port and lever-press responses.

### Experiment 2

In Experiment 1, reward value (magnitude) was manipulated during Pavlovian conditioning. This complicates data interpretation because it conflates the influence that reward value has on conditioned responding with the potential influence that expected reward value may have on the regulation of PIT performance. Consider the finding that rats decrease their rate of lever pressing when presented with a cue that predicts the delivery of 3 or 9 pellets. This may reflect a previously established tendency for such cues to elicit a competing food-port response, which may be performed in an automatic and inflexible manner. Thus, the results of Experiment 1 do not provide definitive support for the hypothesis that reward-predictive cues prompt a flexible shift from lever pressing to checking the food-port based on expected reward value, even though they are generally compatible with this account.

The post-training reward devaluation procedure avoids this issue and provides a more direct test of the degree to which a response is performed flexibly, based on an internal representation of the expected reward (Colwill and Rescorla 1990; Holland 2004; Holland and Straub 1979; Rescorla 1994). We applied this approach to probe the influence of expected reward value on PIT performance using the same basic task used in Experiment 1, but with a few key changes. In Experiment 2, rats were once again trained to press a lever for a grain-based food pellet, after which they were given differential Pavlovian conditioning with two distinct CS-reward contingencies. One CS signaled the delivery of 3 banana pellets and the other CS signaled the delivery of 3 chocolate pellets. Based on the results of Group 3/3 in Experiment 1, we anticipated that this training would establish cues that suppress lever pressing and elicit high levels of food-port activity during the PIT test. However, in the current experiment, we established a conditioned taste aversion to one of the two Pavlovian rewards after training but prior to testing. Thus, while the two CSs signaled equally valuable rewards during training, only one of these CSs signaled a valuable reward at test, with the other signaling a recently devalued reward. According to the cognitive control account, this treatment should selectively disrupt the behavioral effects of the CS associated with the devalued reward since this cue no longer signals a need to suppress lever pressing and check the food-port. However, if the behavioral effects of these CSs are established during training and expressed in a habitual or automatic manner at test, then reward devaluation should have no impact on task performance.

## Methods

### Animals and Apparatus

Sixteen experimentally naïve adult male Long Evans rats (Charles River) were used in Experiment 2. As in Experiment 1, they arrived at the facility at approximately 10 weeks of age. Husbandry, feeding, and experimental conditions were identical to that of Experiment 1. The experiment was conducted in 16 operant chambers identical to the ones used in Experiment 1.

### Procedure

#### Magazine training

Rats received two magazine-training sessions as described in Experiment 1.

#### Instrumental training

Instrumental training was conducted as in Experiment 1 with the exception that all rats received two daily sessions of FR-1 training, which were followed by 1 day of RI-15, 2 days of RI-30, and 10 days of RI-45 training.

#### Pavlovian training

Pavlovian training was conducted according to the same conditions that were used for Experiment 1’s Group 3/3, except that each cue predicted a different type of food pellet. The food pellets used during Pavlovian training also differed from the one used during instrumental training. For half of the rats (*n* = 8) a 3-kHz tone predicted the delivery of three chocolate-flavored, sucrose-based pellets and a 10-Hz clicker predicted the delivery of three banana-flavored, sucrose-based pellets (45 mg; Dustless Precision Pellets, BioServ). The other rats (*n* = 8) received the alternative arrangement (i.e., the tone predicted banana pellets and the click predicted chocolate pellets). As for Experiment 1, Pavlovian training sessions involved 4 pseudorandomly-alternating presentations of each CS (8 total trials per session) separated by 120-s ITI. Pavlovian training took place over 25 sessions.

#### Specific reward devaluation by conditioned taste aversion (CTA)

Following Pavlovian training one of the two food pellets used during Pavlovian conditioning was devalued through conditioned taste aversion learning. One type of food pellet was paired with nausea induced by an injection of lithium chloride (LiCl), whereas the other food pellet was paired with a saline injection (counterbalanced across conditioning contingencies; both reward and stimulus identity). Rats first received 20 g of one type of food pellet in a metal cup in a neutral housing plexiglass cage for 60 min, after which they were given an injection of saline (20 mL/kg) before being returned to the home cage. The following day they received 20 g of the other type of food pellet for 60 min, followed by an injection of 20 mL/kg LiCl (0.15 M) (Balleine and Dickinson 1992; Bouton et al. 2020) before being placed back in their home cage. Rats received a total of three trials (days) with each food-injection arrangement in alternation over six days.

#### Pavlovian-to-instrumental transfer (PIT)

Following CTA, rats received two daily sessions of instrumental retraining (identical to earlier sessions with the RI-45 schedule) followed, on the next day, by a 30-min session of instrumental extinction, as in Experiment 1. On the following day, rats received a PIT test session as in Experiment 1, during which the lever was once again continuously available but inactive. During the test, rats received 4 noncontingent presentations of each 30-s CS in pseudorandom order (ABBABAAB). The ITI was 150 s, and a 5-min interval preceded onset of the first CS. No food pellets were delivered at test.

### Data Analysis

As in Experiment 1, summary measures were obtained from the raw data using MATLAB (The MathWorks; Natick, MA, USA) and analyzed with mixed-effects regression models (Pinheiro and Bates 2000). Even though devaluation had yet to occur, Pavlovian training analyses included fixed effects of group (i.e., banana-vs. chocolate-flavored pellet devaluation) and CS type (i.e., devalued vs. nondevalued) and a by-rat random intercept. Unlike Experiment 1, due to programming error, instrumental-training analysis collapsed across all time bins, and a Wilcoxon rank sum test was used to compare average lever-pressing rates between the rats who were to have banana pellets devalued and those who were to have chocolate pellets devalued. For PIT analyses, because there were no group differences during Pavlovian and instrumental training (as reported below), the fixed-effects structure included CS type and 10-s CS time bin (CS Time; continuous), along with the corresponding two-way interaction (CS Type × CS Time); the random-effects structure included a by-rat random intercept. Similarly, for the analysis of conditioned taste aversion (CTA), which preceded PIT, the mixed-effects model’s fixed-effects structure included the main effects of and interaction between day (1, 2, 3; continuous) and CTA condition (saline, LiCl; categorical), with by-rat random intercepts and slopes as a function of day (restricted to be uncorrelated). For CTA analysis, because the criterion was grams consumed, we employed a Poisson distribution with a log link function (i.e., data were converted to decagrams for analysis). Categorical predictors were effects-coded (i.e., codes sum to 0), and continuous predictors were mean-centered.

As in Experiment 1, our primary measures were lever-press and food-port entry rates. Food-port dwell times were also recorded and analyzed. Outlier detection and removal were also conducted as in Experiment 1. Specifically, data from one rat was removed from instrumental training analyses and data from one rat was removed from Pavlovian training analyses; for PIT analyses, one rat’s lever-press data for trials with the nondevalued CS was removed. Two data points were removed from the CTA analyses. Lastly, 41 of 384 data points (i.e., paired observations of lever-pressing and food-port-entry behavior) were excluded from trial-by-trial analyses of concurrent local changes in both CS-induced lever-pressing and food-port-entry behavior, and 31 of 384 data points (i.e., paired observations of lever-pressing and dwell-time behavior) were excluded from trial-by-trial analyses of concurrent changes in both CS-induced lever-pressing and CS-induced changes in dwell time. As in Experiment 1, the alpha level for all tests was .05, and main effects and interactions are reported as the results of ANOVA *F*-tests.

## Results

### Instrumental and Pavlovian Training

As in Experiment 1, rats were first trained to lever press for food, with the final days of training involving an RI-45-s schedule of reinforcement. While the rats had yet to experience any reward devaluation, we compared the extent to which lever-press rates differed by the eventual groups. Across the final three sessions of instrumental training, mean (*SEM*) lever-press response rates for the rats in which the banana- and chocolate-flavored pellets were later devalued were 15.3 (1.5) and 14.8 (1.4), respectively, which were not significantly different, *p* = .397 (Wilcoxon rank sum test). Likewise, there were no group differences in cue-elicited food-port entry behavior during the final five sessions of Pavlovian training, *F*(1, 27) = 0.01, *p* = .942 [banana devalued: 10.3 (1.3); chocolate devalued: 10.0 (2.1)], and there were no differences in cue-elicited food-port entry behavior between the subsequently devalued versus nondevalued cues, *F*(1, 27) = 1.88, *p* = .182 [devalued: 9.5 (1.9); nondevalued: 10.8 (1.7)].

### Conditioned taste aversion (CTA) training

After initial training, one of the two types of food pellet used during Pavlovian conditioning was devalued through CTA training. As expected, rats selectively avoided consuming the food pellet that was paired with LiCl versus the saline-paired food, *F*(1, 90) = 634.25, *p* < .001 (Figure 5). Analysis also revealed a main effect of day, *F*(1, 90) = 144.18, *p* < .001, and a significant Day × CTA Condition interaction, *F*(1, 90) = 663.63, *p* < .001.

**Figure 5.**
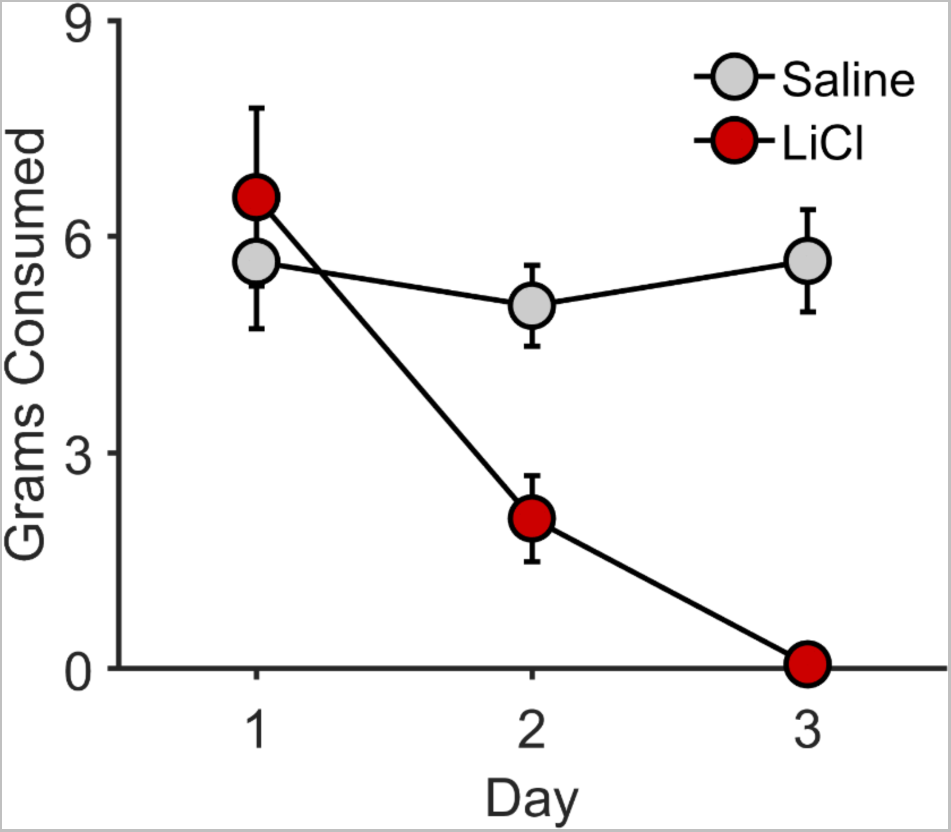
Conditioned taste aversion in Experiment 2. Grams consumed of the food reward paired with saline or lithium chloride (LiCl) as a function of day (trial). Error bars reflect ± 1 between-subjects standard error of the mean.

### Pavlovian-instrumental transfer (PIT)

After CTA training, rats received 2 sessions of instrumental retraining and 1 session of instrumental extinction before undergoing a PIT test. Figure 3 shows cue-elicited changes in lever pressing and food-port entry behavior relative to the 30-s pre-CS period, plotted separately by CS type based on whether or not that cue signaled a devalued reward. For lever pressing (Figure 6A), there was a main effect of CS type, *F*(1, 89) = 6.23, *p* = .014, with the cue that signaled the nondevalued reward suppressing the rate of lever pressing more than the cue that signaled the devalued reward. There was no effect of CS time, *F*(1, 89) = 2.61, *p* = .110, nor was there a CS Time × CS Type interaction, *F*(1, 89) = 0.06, *p* = .809.

**Figure 6.**
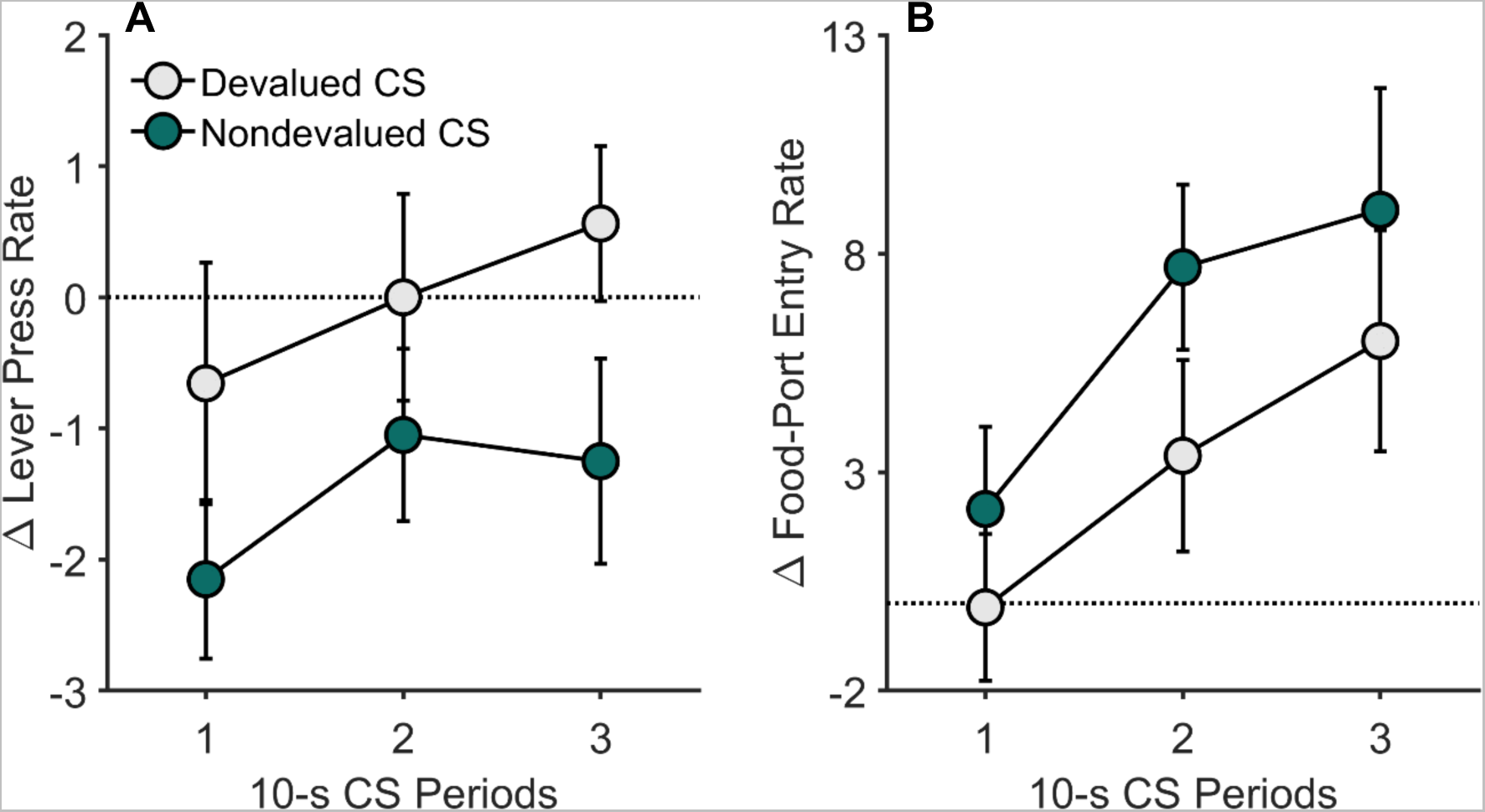
Pavlovian-instrumental transfer in Experiment 2. CS-induced changes in (A) lever-press rates and (B) food-port entries relative to the pre-CS baseline on the Pavlovian-instrumental transfer (PIT) test. These data are plotted separately according to CS Type (Devalued vs. Nondevalued) and show changes in responding over time (10-sec bins) during 30-sec CS presentations (averaged across trials). CS = conditioned stimulus. Error bars reflect ± 1 between-subjects standard error of the mean.

In addition to being less effective in suppressing instrumental behavior, the cue that signaled the devalued reward was less effective in increasing food-port entries versus the cue that signaled the nondevalued reward, *F*(1, 92) = 4.83, *p* = .031 (Figure 6B). There was a main effect of CS Time on cue-induced food-port entries, *F*(1, 92) = 13.25, *p* < .001, but not a CS Time × CS Type interaction, *F*(1, 92) = 0.04, *p* = .833.

As in Experiment 1, such findings raise the possibility that food-port entry behavior may have interfered with lever-press performance during PIT trials through response competition. However, we once again found that local changes in entry rate did not negatively correlate with changes in press rate (Figure 7), as would be expected if cue-elicited food-port behavior prevented rats from pressing the lever. Except for one rat with respect to the nondevalued CS, *r* = .59, *p* = .045, there were no significant within-subject associations between changes in press rate and changes in food-port behavior, *p*’s ≥ .055 (median *p*-value = .522). Moreover, inspection of the peri-event histograms (Figure 8) indicates that these responses again produced only very brief periods of overt competition, which was offset by their coordinated performance as lever-pressàfood-port entry sequences.

**Figure 7.**
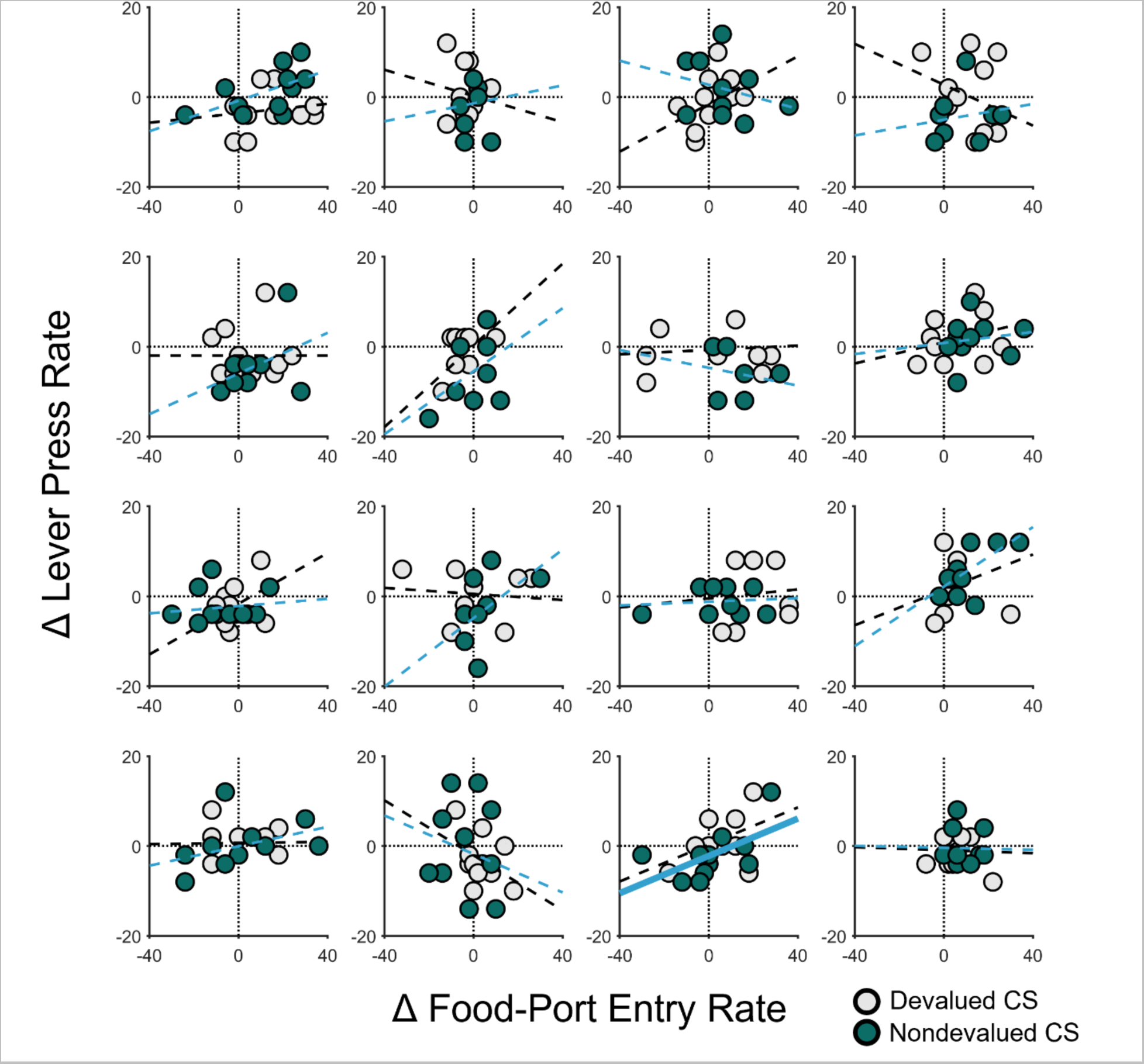
Trial-by-trial relationship between food-port-entry and lever-press rate for each 10-s CS bin during the Pavlovian-instrumental transfer (PIT) test for individual rats in Experiment 2. Scatterplots are presented for each rat. Data are separated by trial (unlike in Figure 6) and 10-s CS bin (as in Figure 6). The abscissa shows the CS-induced changes in food-port-entry rate for individual time bins; the ordinate shows the CS-induced changes in lever-press rate for corresponding time bins. For each CS type for each rat, simple regression lines were fit, with dashed lines indicating that the fit was not significant and bolded lines indicating significance at *p* < .05. CS = conditioned stimulus.

**Figure 8.**
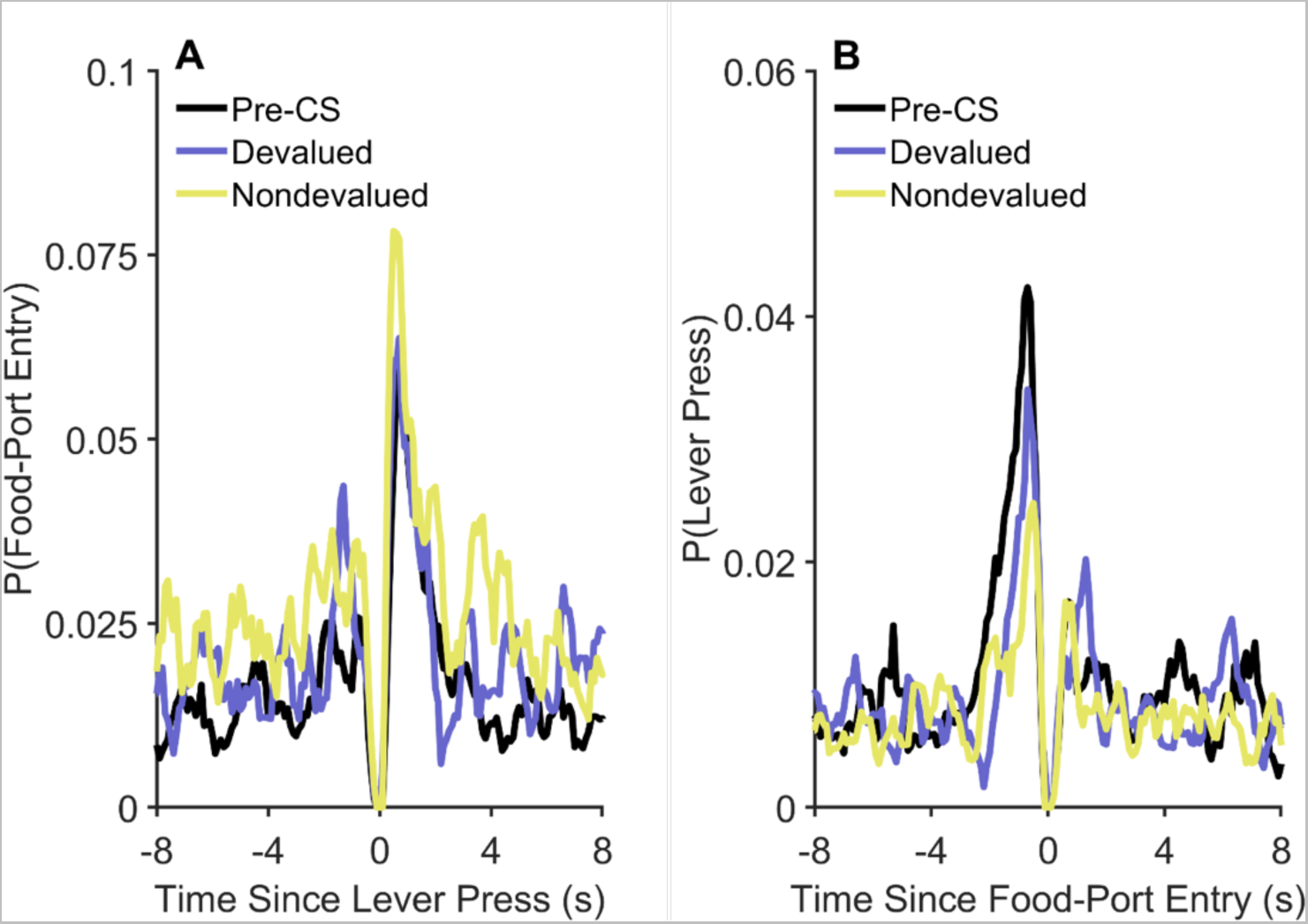
Peri-event histograms for lever-pressing and food-port-entry behavior during the Pavlovian-instrumental transfer test in Experiment 2. (A) Probability of food-port entries as a function of time from each lever press. (B) Probability of lever presses as a function of time from each food-port entry. Note the y-axes are different for Panels A and B.

**Figure 9.**
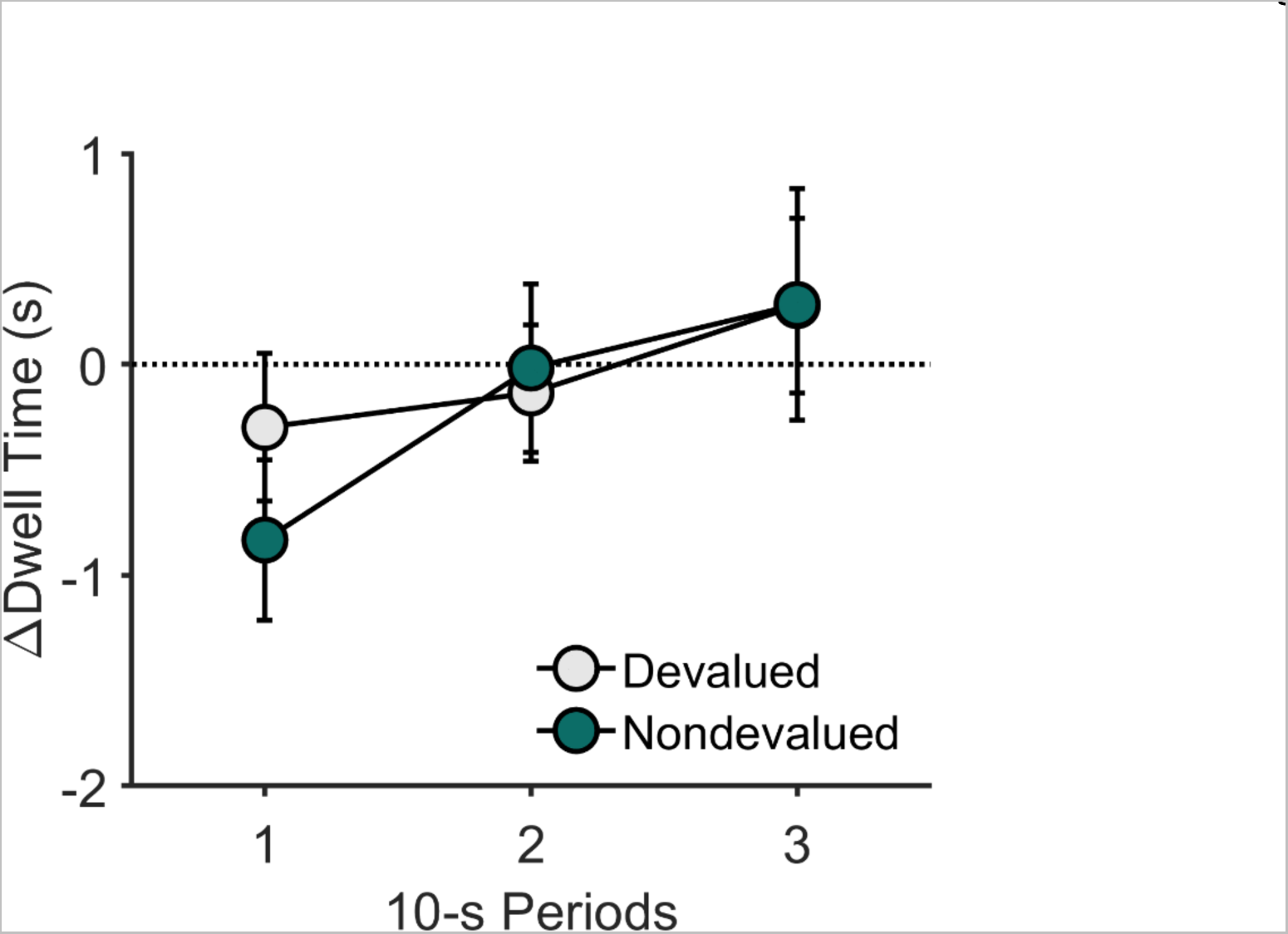
Food-port dwell time (s) during the Pavlovian-instrumental transfer test in Experiment 2. CS-induced changes in the amount of time rats spent in the food-port relative to the pre-CS baseline during the Pavlovian-instrumental transfer (PIT) test. CS = conditioned stimulus. Error bars reflect ± 1 between-subjects standard error of the mean.

**Figure 10.**
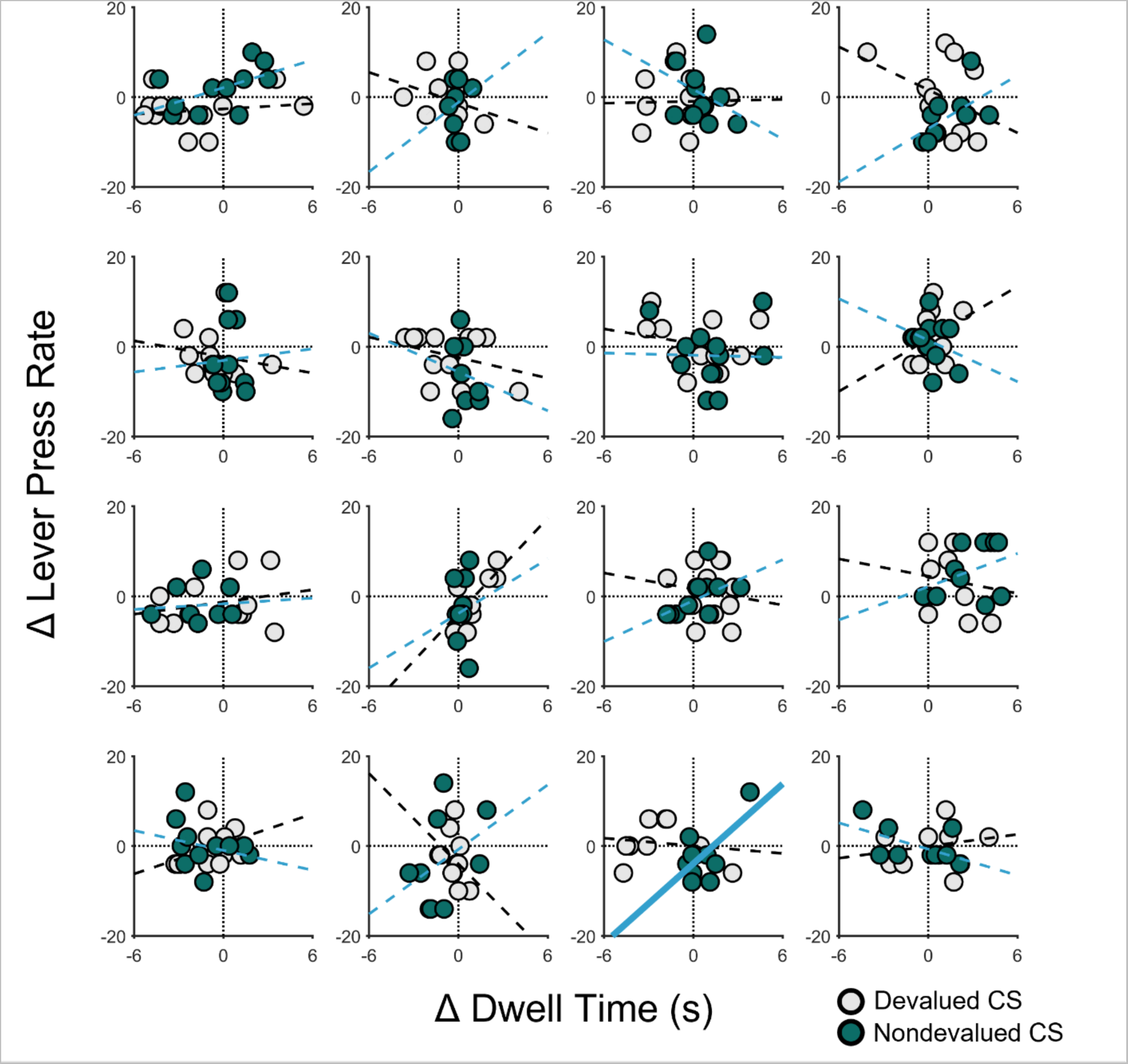
Trial-by-trial relationship between food-port dwell times and lever-press rate for each 10-s CS bin during the Pavlovian-instrumental transfer (PIT) test for individual rats in Experiment 2. Scatterplots are presented for each rat. Data are separated by trial (unlike in Figures 6 and 9) and 10-s CS bin (as in Figures 6 and 9). The abscissa shows the CS-induced changes in dwell time for individual time bins; the ordinate shows CS-induced changes in lever-press rate for corresponding time bins. For each CS type for each rat, simple regression lines were fit, with dashed lines indicating that the fit was not significant and bolded lines indicating significance at *p* < .05. CS = conditioned stimulus.

We also analyzed food-port dwell times (Figure 9) during the PIT test, which revealed that rats waited with their heads in the food-port to a similar degree during both CSs. Specifically, while dwell time increased over time during CS presentations, *F*(1, 92) = 8.55, *p* = .004, there was no effect of CS type, *F*(1, 92) = 0.34, *p* = .564, nor a CS Time × CS Type interaction, *F*(1, 92) = 0.87, *p* = .354. Furthermore, except for one rat with respect to the nondevalued CS, *r* = .66, *p* = .027, we found no within-subject correlations between CS-induced changes in dwell time and CS-induced changes in lever pressing, *p*’s ≥ .050 (median *p*-value = .440; Figure 10). Therefore, it seems unlikely that the effect of reward devaluation on cue-elicited pressing was mediated by interference with food-port behavior.

## Discussion

These results show that Pavlovian reward-predictive cues can suppress instrumental behavior and increase food-port activity in a flexible manner that depends on the current value of the expected reward. It is worth noting that the devalued CS had little influence on instrumental behavior in either direction. It is possible that cues which normally suppress behavior may acquire latent motivational properties that can be expressed under certain circumstances. For instance, a cue that signals a devalued reward might become an effective motivator of instrumental behavior if the devaluation treatment removes the inhibitory influence of reward expectancy without disrupting that cue’s motivational properties. As noted above, previous studies have shown that reward devaluation typically does not disrupt the motivational influence of reward-paired cues (Colwill and Rescorla 1990; Holland 2004; Rescorla 1994). However, such studies have explored a limited range of parameters, typically involving cues that are loosely correlated with reward delivery. Thus, it is possible that cues that signal precise information about the time of reward delivery, such as the CSs used here, acquire reward-specific motivational properties that are gated by expected reward value. Interestingly, (Colwill and Rescorla 1990) found that while cues generally retain their ability to motivate instrumental behavior following reward devaluation, this was not the case for cues that signal forced intra-oral reward delivery, suggesting that Pavlovian incentive motivation is indeed modulated by reward value under certain circumstances.

It is also possible that highly predictive cues, such as our CSs, do not acquire significant motivational properties, or at least do not acquire properties that allow them to motivate instrumental behavior. It is also notable that the current experiment used a “general” PIT design, in that different rewards were used during instrumental and Pavlovian conditioning phases, which may have limited the excitatory motivational influence of the CSs. In contrast, in Experiment 1 and related studies (Marshall and Ostlund 2018), the same reward was used during both conditioning phases, which may support a stronger excitatory PIT effect. This procedural difference may also explain why, in Experiment 2, we did not observe an initial rise or progressive decrease in lever pressing during CS presentations, as in Experiment 1 and previous studies (Marshall and Ostlund 2018). Yet another possibility is that the devaluation procedure used here was incomplete or for some other reason failed to completely disrupt the suppressive influence of reward expectancy on instrumental behavior. In this case, the motivational potential of the devalued CS would remain unexpressed. While future studies will be needed to assess these possibilities, the current findings provide strong evidence that the suppressive influence of reward-predictive cues is flexibly modulated by expected reward value.

### General Discussion

The current study shows that reward expectancy plays an important role in regulating the expression of PIT performance. In Experiment 1, we found that cues signaling a relatively small reward (1 pellet) elicit an immediate and sustained increase in lever pressing while only modestly increasing food-port activity. In contrast, cues signaling either intermediate (3 pellets) or large (9 pellets) magnitude rewards suppressed lever pressing, particularly at the end of the cue period, near the expected time of reward delivery. Experiment 2 provided more definitive evidence that reward-predictive cues suppress instrumental behavior in a flexible, goal-directed manner. Specifically, we found that the tendency for rats to withhold lever pressing when presented with such a cue was attenuated if that cue signaled a devalued reward.

While the current findings show that expected reward value governs the suppressive influence of reward-predictive cues on instrumental behavior, this is usually not the case for the excitatory motivational influence of cues (i.e., Pavlovian incentive motivation). As noted earlier, cues that are effective in motivating instrumental behavior tend to be long in duration (e.g., 2-min) and signal sporadic reward delivery (Crombag et al. 2008; Lovibond 1981). Previous studies have shown that the motivational influence of such cues is surprisingly insensitive to post-training reward devaluation (Colwill and Rescorla 1990; Holland 2004; Rescorla 1994). This has been shown for both the nonspecific, response-invigorating effect of such cues (Holland 2004), referred to as general PIT, as well as the specific form of PIT, in which cues selectively increase the performance of instrumental actions with which they share a common outcome (Colwill and Rescorla 1990; Holland 2004; Lingawi et al. 2022; Rescorla 1994; Sommer et al. 2022) (but see, Panayi and Killcross 2022). Thus, the excitatory PIT effect appears to be driven by a Pavlovian motivational process which compels instrumental behavior in an automatic or habitual manner, as opposed to prompting a more deliberative process of goal-directed decision making. This conclusion is bolstered by reports that instrumental training protocols that promote habit formation also render performance more sensitive to Pavlovian incentive motivation (Holland 2004; Wiltgen et al. 2012).

We propose that although Pavlovian incentive motivation is normally deployed in an automatic manner, it is susceptible to top-down cognitive control under certain conditions, allowing it to be flexibly regulated based on the current value of an expected reward. Reward timing appears to be another important factor involved in regulating Pavlovian incentive motivation. For instance, lever pressing is suppressed by cues signaling imminent reward and stimulated by cues signaling long delays in reward delivery (Crombag et al. 2008; Lovibond 1981; Marshall and Ostlund 2018). Moreover, we have shown that cues signaling relatively short, predictable delays in reward delivery provoke a gradual decrease in instrumental behavior that is most pronounced at the expected reward delivery time (Marshall and Ostlund 2018), a trend that was also apparent in Experiment 1. Pavlovian incentive motivation also appears to be negatively regulated by expected reward probability. For instance, we have shown that cues signaling that a low probability of reward tend to increase instrumental behavior, whereas cues signaling a high probability of reward tend to decrease instrumental behavior (Marshall et al. 2020). In addition to signaling information about reward likelihood, timing, magnitude, and incentive value, cues also shape instrumental behavior by signaling qualitative details about the flavor and texture of expected rewards, as is evident in the specific PIT effect (Balleine 2016; Balleine and Ostlund 2007; Delamater 2011). This ability for cues to bias action selection to promote the pursuit of expected rewards is dissociable from their more general motivational effects (Corbit and Balleine 2016; Ostlund and Maidment 2012).

Our conceptual framework is generally compatible with behavior systems theory (Timberlake et al. 1982), which assumes that cues shape conditioned behavior by relaying information about the timing and location of food. According to this account, cues that signal uncertain or delayed reward trigger a general search mode (e.g., sign-tracking), whereas cues signaling imminent reward trigger a focal search mode (e.g., goal-tracking). Research on the influence of expected reward probability (Anselme et al. 2013; Davey and Cleland 1984) and reward timing (Timberlake et al. 1982) on Pavlovian conditioned responding largely support these predictions. By extension, this theory also readily explains the influence of these factors on PIT performance if one makes the straightforward assumption that instrumental behavior is a general search activity and food-port entry is a focal search activity. However, the behavior systems account does not specify how rewards are represented during learning and therefore does not make useful predictions about whether treatments such as reward devaluation should differentially affect general versus focal search behavior.

Our framework is more directly aligned with Konorski’s theory (1967, p. 276-280) that Pavlovian conditioning involves independent motivational and cognitive learning processes. According to this model, reward-paired cues rapidly become associated with the emotional-motivational properties of reward, allowing them to elicit preparatory conditioned responses and instrumental behavior. This motivational capacity extends to all reward-paired cues, regardless of their temporal relationship with reward. However, short cues that signal imminent reward delivery are assumed to be unique in their tendency to form a separate association with a sensorily detailed representation of reward. Activating this representation results in what is essentially a cognitive reward expectancy, which is assumed to both evoke reward-specific consummatory behaviors and inhibit concurrent activity in the emotional-motivational pathway, thereby suppressing the impulse to act. This model readily accounts for the opposing effects of reward imminence and reward probability on lever pressing and food-port approach behavior during PIT, but also accounts for the opposing effects of reward magnitude and incentive value on these behaviors, at least if one assumes that activation of the sensory reward representation is greater when the predicted reward is large and desirable than when it is small or undesirable. However, it should be noted that Konorski’s account does not readily explain the specific PIT effect, since it assumes that the process of activating an outcome-specific reward representation should also dampen a cue’s motivational effects. Even more problematic, Delamater and Holland (2008) found that cues that signal imminent reward (2-sec delay) are less effective at eliciting specific PIT than cues signaling longer delays in reward delivery (20- to 180-sec). Such findings are at odds with Konorski’s account and indicate that temporal and sensory features of reward are differentially encoded during Pavlovian conditioning and exert distinct effects on instrumental performance.

We suggest that this inhibitory influence of reward-predictive cues represents a form of top-down cognitive control that allows for flexible and efficient foraging behavior. Seeking out new rewards through exploratory instrumental behavior is useful when rewards are scarce, unpredictable, or difficult to obtain. However, when reward delivery is imminent, such behavior becomes wasteful and may interfere with reward retrieval and consumption. We suggest that rapidly shifting from reward seeking to reward retrieval is particularly advantageous when a highly desirable reward is expected, because there is more to lose in such situations. Conversely, there is less need to avoid reward seeking when a small or devalued reward is expected since the potential loss is relatively low. In this case, it may be more appropriate to delay searching for the expected reward, perhaps until ongoing instrumental behavior has been completed.

While the current paper focuses on how Pavlovian learning influences instrumental behavior, similar processes appear to shape the expression of conditioned approach behavior. Rats’ tendency to sign-track (i.e., approach the cue) is commonly assumed to reflect a generalized motivational response (incentive salience), whereas goal-tracking (i.e., checking the food-port) is assumed to be mediated by a cognitive expectancy of reward (Flagel et al. 2009; Robinson et al. 2018; Sarter and Phillips 2018). This distinction is supported in part by findings that goal-tracking tends to be more sensitive than sign-tracking to reward devaluation (Amaya et al. 2020; but see Derman et al. 2018; Keefer et al. 2020; Morrison et al. 2015; Patitucci et al. 2016). There is now a large body of studies that have adopted this framework to elucidate the neural mechanisms of motivation and cognitive control and explore how individual variability in conditioned approach behavior relate to addiction vulnerability (Anselme and Robinson 2020; Flagel et al. 2009; Robinson et al. 2018; Sarter and Phillips 2018). We suggest that the PIT paradigm provides a complementary approach that more directly models cognitive control over voluntary instrumental reward-seeking behavior.

At first glance, the current findings appear to be incompatible with current theories of incentive learning (Dayan and Balleine 2002; McClure et al. 2003; Zhang et al. 2009) which assume that the motivational properties of reward-predictive cues are directly linked to their state values – i.e., the total delay-discount reward expected based on a cue, as determined by temporal difference learning. These theories make the straightforward and intuitive prediction that cues signaling that an upcoming reward will be large and desirable should be more motivating than cues that predict a small or undesirable reward. In contrast, the current study found evidence of the opposite relationship; that is, the motivational effects of a cue varied inversely with the value of the predicted reward. This theoretical approach is also difficult to reconcile with previous findings that the motivational effects of a cue are negatively regulated by reward probability (Marshall et al. 2020) and reward imminence or timing (Marshall and Ostlund 2018).

However, this interpretation assumes that the degree to which a cue stimulates instrumental performance provides a reliable and selective readout of its incentive motivational properties. While this is indeed the central assumption of the PIT paradigm (Corbit and Balleine 2016; Holmes et al. 2010; Rescorla and Solomon 1967), it is possible that a cue’s motivational properties can also stimulate other appetitive behaviors such as food-port activity. If this were the case, it would be a mistake to conclude that cues that signal large, valuable reward lack motivational properties, since they are generally effective in eliciting some form of appetitive behavior, if not instrumental performance *per se*. It is also possible that cues are indeed assigned motivational properties in line with their state values, as has been posited (Dayan and Balleine 2002; McClure et al. 2003; Zhang et al. 2009), but that these properties remain latent under certain conditions, such as when they are actively suppressed to allow for other more controlled reward-seeking actions or strategies, such as devaluation-sensitive food-port entry behavior. This is in line with previous studies showing that cues which signal imminent delivery of a valuable reward can motivate instrumental behavior if the predicted reward does not require the production of an conflicting consummatory response (Baxter and Zamble 1982; LeBlanc et al. 2012; Lovibond 1983), or if the conflicting behavior (e.g., food-port activity) has been extinguished prior to testing (Holmes et al. 2010; Marshall and Ostlund 2018). While such findings are typically interpreted in terms of overt response competition, we suggest that they may instead reflect conditions under which it is not advantageous to suppress Pavlovian incentive motivation or its excitatory influence on instrumental reward-seeking behavior. Regardless of which of these accounts is more accurate, both are at least compatible with the assumption that the motivational properties of a cue are linked to its state value (Dayan and Balleine 2002; McClure et al. 2003; Zhang et al. 2009).

We have proposed (Ostlund and Marshall 2021) that this tendency for reward-predictive cues to suppress instrumental behavior (positive conditioned suppression), may be useful for assaying cognitive control over cue-motivated behavior and how it can go awry to produce maladaptive behavior. For instance, we have shown that adolescent male rats are impaired in using expected reward probability to shift from instrumental behavior to food-port entry during PIT (Marshall et al. 2020). Whereas these rats increased their instrumental behavior when presented with a cue that signaled a high probability of reward, a control group of adult male rats showed the opposite effect, suppressing their instrumental behavior while focusing their activity at the food-port. Interestingly, cues that signaled a low probability of reward elicited a similar increase in instrumental behavior in both adolescent and adult groups, which indicates that adolescent rats were not simply more motivated by reward-paired cues. Using a similar approach, we have shown that rats with a history of repeated cocaine exposure are impaired in regulating their cue-motivated behavior, in this case based on the expected time of reward delivery (Marshall and Ostlund 2018).

The current study suggests that such deficits reflect a failure of cognitive control over cue-motivated behavior, which is normally expressed in a flexible and goal-directed manner. Further work is needed to validate this framework. For instance, more should be done to determine if cues normally motivate instrumental behavior through an automatic process (that is, without considering the current value of the predicted reward) and whether this depends on Pavlovian conditioning parameters such as CS-reward interval. More should also be done to characterize the nature of the positive conditioned suppression effect. While our findings suggest that this effect is not a simple product of response competition, it would be a mistake to conclude that response conflict plays no role in this phenomenon. Indeed, theories of cognitive control assume that one of its core functions is to flexibly resolve conflict between competing response tendencies (Botvinick and Braver 2015; Braver 2012). Although some findings suggest that positive conditioned suppression may be limited to situations that involve conflict between overt motor responses (Baxter and Zamble 1982; Lovibond 1983), this question deserves more attention. For instance, previous reports suggest that positive conditioned suppression is particularly pronounced when instrumental behavior is reinforced on a high-effort ratio schedule (Kelly 1973; Lovibond 1981; Soltysik et al. 1976), which suggests that this effect may reflect an adaptive response to reduce unnecessary energy expenditure.

## Acknowledgements

The authors are supported by NIH grants DA046667 (SBO), MH12685 (SBO), MH106972 (SBO), DA050116 (BH). Raw data and code are available from the corresponding author upon request. This article was published as a preprint on bioRxiv: doi: https://doi.org/10.1101/2021.04.08.438512.

